# Kinesin-1 trans-synaptically regulates synaptic localization of SARM1 for asymmetric neuron diversification

**DOI:** 10.64898/2026.01.05.697830

**Authors:** Anaam Khalid, Peter Sahyouni, Jun Yang, Shengyao Yuan, Rui Xiong, Chiou-Fen Chuang

## Abstract

The *Caenorhabditis elegans* AWC olfactory neuron pair differentiates stochastically into two distinct subtypes, default AWC^OFF^ and induced AWC^ON^. A calcium signaling complex assembled by the TIR-1/SARM1 adaptor protein is transported from the AWC cell body to the axons, where it cell autonomously specifies the AWC^OFF^ subtype through lateral signaling. UNC-104, the *C. elegans* homolog of the kinesin-3 motor protein KIF1A, acts non-cell autonomously in AWC^ON^ to control the synaptic localization of the TIR-1 signaling complex in promoting AWC^OFF^. Here, we identify a non-cell-autonomous role of *unc-116/kinesin-1*, similar to that of *unc-104/kinesin-3*, in promoting AWC^OFF^. *unc-116* mutants, similar to *unc-104* mutants, enhance the 2AWC^ON^ phenotype of a hypomorphic *tir-1* mutant. Overexpression of *unc-116* in AWC causes a 2AWC^OFF^ phenotype, the same as the *tir-1* overexpression phenotype. Like UNC-104, UNC-116 plays a non-cell-autonomous role in the AWC^ON^ cell to promote AWC^OFF^ cell subtype by regulating the dynamic trafficking of TIR-1 along the AWC axon. UNC-116 is strikingly colocalized with UNC-104, while both are generally adjacent to TIR-1. Taken together, these results suggest a model in which UNC-116/kinesin-1 and UNC-104/kinesin-3 may work cooperatively to transport some unknown presynaptic factor(s) in the future AWC^ON^ cell that trans-synaptically regulates the dynamic trafficking of the TIR-1/SARM1 signaling complex to postsynaptic regions of the AWC axons in promoting the AWC^OFF^ subtype.

## Introduction

Microtubule molecular motor proteins, kinesin and dynein, play a critical role in development by facilitating the intracellular transport of cellular components essential for cell division, neuronal development, and morphogenesis. Dyneins are exclusive to retrograde (minus-end-directed) transport (Hou and Witman 2015; Viswanadha *et al*. 2017; Reck-Peterson *et al*. 2018), while kinesins comprise a larger superfamily responsible for anterograde (plus-end-directed) transport and microtubule structure formation and stability (Siddiqui 2002; Miki *et al*. 2005; Hirokawa and Noda 2008; Ali and Yang 2020). Kinesins, in addition to their cell-autonomous functions, also play a role in non-cell-autonomous processes, where kinesins in one cell can regulate cellular activity in neighboring cells through neuronal signaling (Chang *et al*. 2011; Auer *et al*. 2015; Glomb *et al*. 2023; Quesnelle *et al*. 2023). For example, the *C. elegans* KIF1A homolog UNC-104/kinesin-3 functions non-cell autonomously in one AWC olfactory neuron subtype to regulate the dynamic transport of TIR-1/SARM1 along AWC axons in the contralateral AWC neuron to specify the other AWC subtype through lateral signaling (Chang *et al*. 2011).

The *C. elegans* AWC olfactory neuron pair uses calcium-dependent lateral signaling to differentiate into two subtypes, AWC^OFF^ and AWC^ON^, in a stochastic manner. The default AWC^OFF^ expresses the G protein-coupled receptor (GPCR) gene *srsx-3* and senses the odor pentanedione, while the induced AWC^ON^ expresses the GPCR gene *str-2* and senses the odor butanone (Troemel *et al*. 1999b). Wild-type worms display the 1AWC^ON^/1AWC^OFF^ phenotype, with one AWC^ON^ and one AWC^OFF^ neuron. Mutations that disrupt the function or regulation of the calcium signaling pathway genes can result in the loss of one of these subtypes, causing a 2AWC^ON^ or 2AWC^OFF^ mutant phenotype.

The default AWC^OFF^ subtype is specified by a voltage-gated calcium channel (UNC-2, EGL-19, UNC-36) and a downstream calcium-dependent MAP kinase signaling pathway. In this pathway, the TIR-1/SARM1 adaptor protein assembles a calcium signaling complex that consists of the upstream UNC-43 calcium/calmodulin-dependent protein kinase (CaMKII) and the downstream MAP kinase cascade (Troemel *et al*. 1999b; Sagasti *et al*. 2001; Tanaka-Hino *et al*. 2002; Chuang and Bargmann 2005; Bauer Huang *et al*. 2007). The TIR-1 signaling complex is transported from the AWC cell body to postsynaptic sites in the AWC axon, where the two AWC cells communicate to specify the AWC^OFF^ subtype (Chuang and Bargmann 2005; Chang *et al*. 2011; HSIEH *et al*. 2014). The induced AWC^ON^ subtype is specified by intercellular calcium signaling through a transient NSY-5 gap junction neural network, the NSY-4 claudin adhesion protein, and the downstream SLO BK potassium channels, all of which antagonize the calcium signaling pathway (Vanhoven *et al*. 2006; Chuang *et al*. 2007; Schumacher *et al*. 2012; Alqadah *et al*. 2016).

TIR-1 acts cell autonomously to specify the AWC^OFF^ subtype (Chuang and Bargmann 2005), while UNC-104/kinesin-3, a homolog of KIF1A, functions trans-synaptically in the AWC^ON^ cell to regulate the dynamic transport of TIR-1 along AWC axons in promoting the AWC^OFF^ subtype (Chang *et al*. 2011). However, the non-cell-autonomous mechanism regulating the axonal transport of TIR-1 in the AWC^OFF^ specification has yet to be elucidated.

Here, we identify a non-cell-autonomous role of UNC-116/kinesin-1, similar to that of UNC-104/kinesin-3, in the synaptic localization of TIR-1 for the AWC^OFF^ specification. Our results show that *unc-116* mutations, like those of *unc-104*, increase the penetrance of the 2AWC^ON^ phenotype in a hypomorphic *tir-1* mutant background. In addition, *unc-116* overexpression, similar to *tir-1* overexpression, causes a 2AWC^OFF^ phenotype. Our genetic data suggest that *unc-116*, like *unc-104*, acts non-cell autonomously in the AWC^ON^ cell to promote the AWC^OFF^ subtype. Moreover, dynamic TIR-1 trafficking along AWC axons is regulated by *unc-116*, similar to *unc-104*. Consistent with our genetic data, UNC-116, like UNC-104, is localized adjacent to TIR-1 and colocalizes with UNC-104 in AWC axons. Together, our results suggest that UNC-116/kinesin-1, alongside UNC-104/kinesin-3, may transport some unknown presynaptic factor(s) in the pre-AWC^ON^ cell that trans-synaptically regulates trafficking of the TIR-1 signaling complex in AWC axons to promote the specification of the AWC^OFF^ subtype.

## Results

### *unc-116/kinesin-1* genetically interacts with *tir-1/SARM1* in promoting the AWC^OFF^ subtype

Previous studies have shown that UNC-104/kinesin-3 is required for the localization of TIR-1/SARM1 at the postsynaptic regions of AWC axons to promote the AWC^OFF^ subtype (Chang *et al*. 2011). To determine whether UNC-116 (the sole heavy chain kinesin-1 in *C. elegans*), like UNC-104/kinesin-3, plays a role in the axonal transport of TIR-1 for the AWC^OFF^ subtype specification, we first analyzed AWC asymmetry phenotypes in single and double mutants of *unc-116* and *tir-1*.

Because *unc-116* null alleles are lethal (Meyerzon *et al*. 2009), other viable *unc-116* mutant alleles were used for the analysis of AWC asymmetry phenotypes. The *unc-116(e2310)* reduction-of-function allele contains a Tc5 transposon insertion after residue 692 in the stalk domain (Patel *et al*. 1993). The *unc-116(rh24)* allele is a gain-of-function allele that contains two missense mutations in the motor domain (I304M and E338K) (Patel *et al*. 1993). The *unc-116(rh24sb79)* allele, a reduction-of-function revertant from the *rh24* background, contains one additional missense mutation in the motor domain (Yang *et al*. 2005). All three *unc-116* mutant alleles, like *unc-104* mutants, displayed wild-type AWC asymmetry in 97-100% of animals (Figure 1A and 1B, rows 1-4, and 12). However, *tir-1(ky388ts); unc-116(e2310)* double mutants displayed a significant increase in the 2AWC^ON^ phenotype of *tir-1(ky388ts)* mutants from 58% to 100% at 20°C (Figure 1A and 1B, rows 6-7) and 18% to 84% at 15°C (Figure 1A and 1B, rows 9-10). Similar results were observed in *tir-1(ky388ts); unc-104(e1265)* double mutants (Chang *et al*. 2011). Together, these results suggest that *unc-116/kinesin-1*, like *unc-104/kinesin-3* (Chang *et al*. 2011), genetically interacts with *tir-1/SARM1* to promote the AWC^OFF^ subtype.

**Figure 1.**
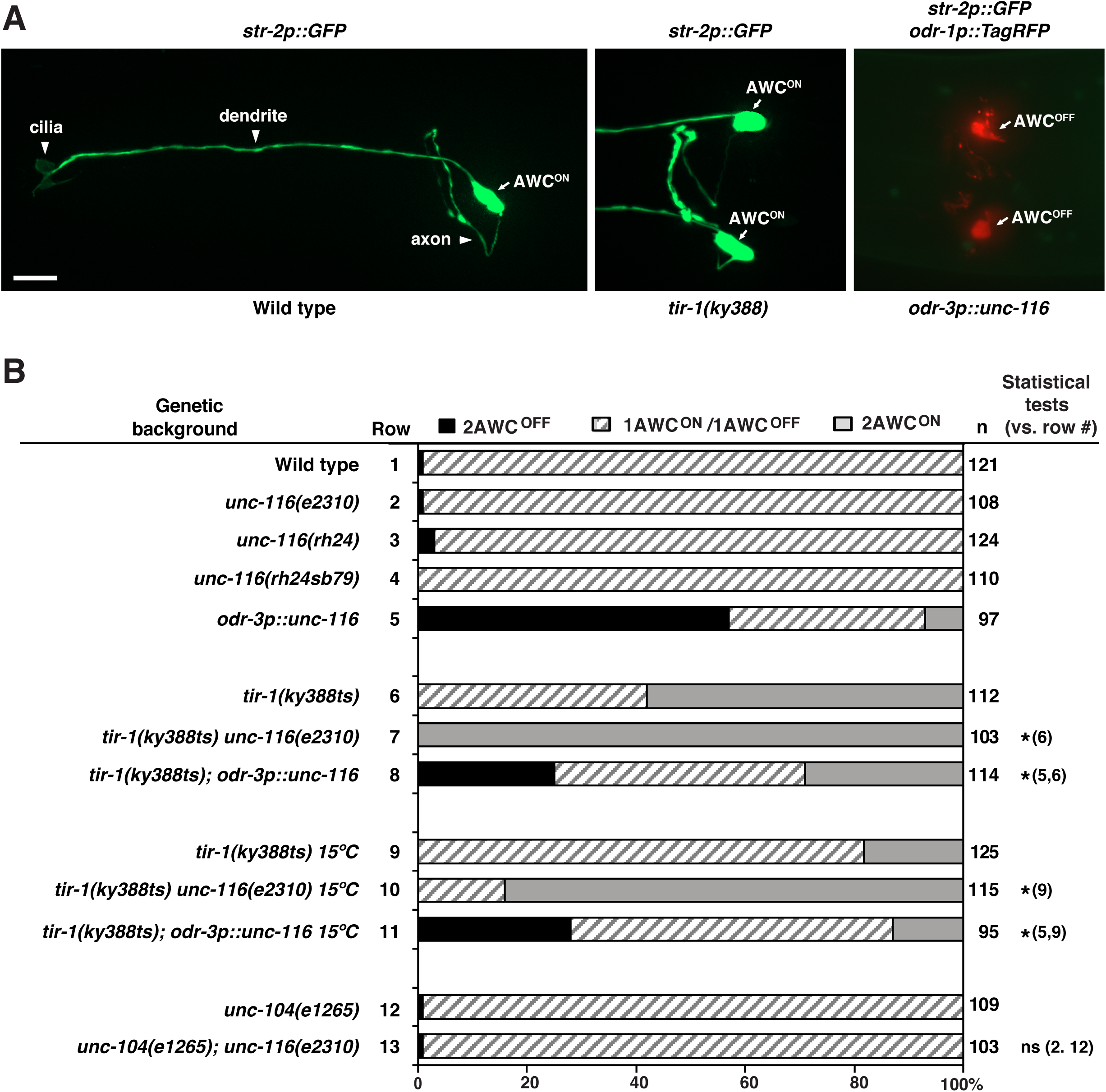
*unc-116/kinesin-1* genetically interacts with *tir-1/SARM1* to promote the AWC^OFF^ cell subtype. **(A)** Images of wild type and *tir-1(ky388)* mutants, expressing *str-2p::GFP* (AWC^ON^ marker), and *odr-3p::unc-116* transgenic animals, expressing *str-2p::GFP* and *odr-1p::TagRFP* (general AWC marker), in the adult stage. Wild-type animals display a 1AWC^ON^/1AWC^OFF^ phenotype. The AWC^OFF^ cell is lost in *tir-1(ky388)* mutants, resulting in a 2AWC^ON^ phenotype, and the AWC^ON^ cell is lost in *odr-3p::unc-116* animals, resulting in a 2AWC^OFF^ phenotype. Arrows indicate the AWC cell body. Scale bar, 10μm. Anterior to the left and ventral at the bottom. **(B)** Expression of the AWC^ON^ marker *str-2p::GFP* in adult wild type, mutants, and transgenic lines. *n,* total number of animals scored. Animals were grown at 20℃, unless otherwise indicated. A *Z*-test was used to make statistical comparisons. Asterisks indicate comparisons that are significant at *p* < 0.5. ns, not significant.

In addition, overexpression of *unc-116* in AWC, from the *odr-3p::unc-116* transgene, causes a 2AWC^OFF^ phenotype (Figure 1B, row 5), similar to the *tir-1* overexpression phenotype (Chuang and Bargmann 2005). Overexpression of *unc-116* partially and significantly suppressed the *tir-1(ky388ts)* 2AWC^ON^ phenotype at 20°C and 15°C (Figure 1B, rows 6, 8, 9, and 11). These results are consistent with the role of *unc-116* as well as the genetic interaction between *unc-116* and *tir-1* in promoting AWC^OFF^.

Both *unc-116(e2310)* and *unc-104(e1265)* mutants displayed wild-type AWC asymmetry (Figure 1B, rows 2 and 12). In order to determine whether this result was due to a functional redundancy between *unc-116* and *unc-104*, we analyzed AWC asymmetry phenotypes in *unc-104(e1265); unc-116(e2310)* double mutants. *unc-104(e1265)* is a partial loss-of-function allele containing the amino acid substitution D1497N in the PH domain, resulting in reduced PI(4,5)P(2) binding (Byrd *et al*. 2022). We found that *unc-104(e1265); unc-116(e2310)* double mutants did not display any significant defects in AWC asymmetry (Figure 1B, row 13). These results suggest the possibility that other, unidentified motor proteins may have overlapping roles with UNC-104/kinesin-3 and UNC-116/kinesin-1 in AWC asymmetry.

### UNC-116/kinesin-1 is localized in AWC cell bodies and axons

It was previously shown that *unc-116* was broadly detected in most neurons, muscle, and pharynx tissue (Sakamoto *et al*. 2005). To determine whether *unc-116* is expressed in AWC cells, an integrated transgene *unc-116::GFP* (McNally *et al*. 2010), expressing the UNC-116::GFP fusion protein driven by the *unc-116* promoter, was co-expressed with the AWC marker transgene *odr-1p::TagRFP.* UNC-116::GFP was detected in the cell bodies of many head neurons, including both AWC cells, and the nerve ring (where axons of head neurons are located) at the first larval stage (Figure 2A). To further determine whether UNC-116 is localized in AWC axons, UNC-116::GFP was expressed in AWC cells using the *odr-3* promoter. In addition to AWC cell bodies, UNC-116::GFP was localized in a punctate pattern along AWC axons (Figure 2B).

**Figure 2.**
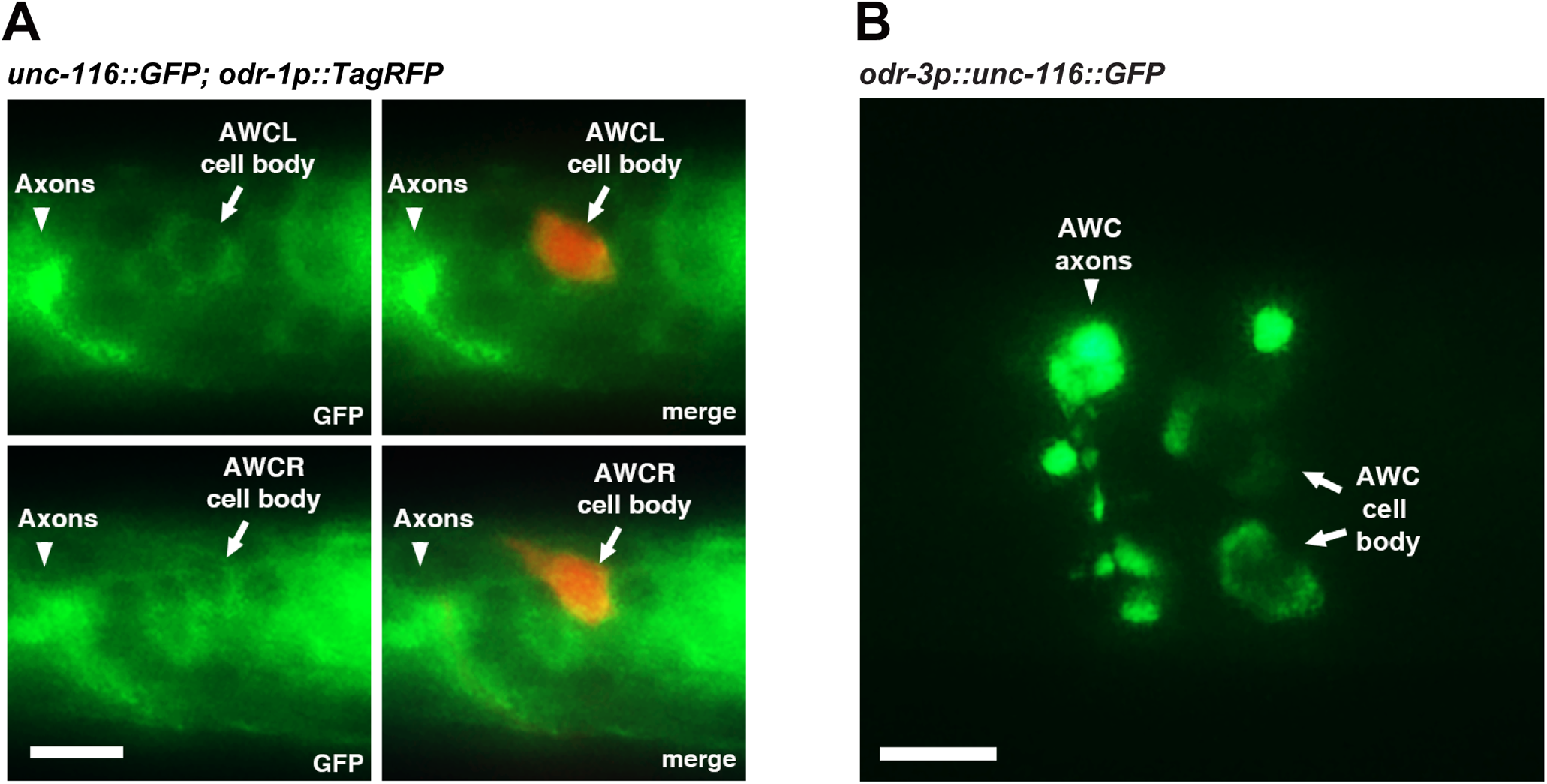
UNC-116/kinesin-1 is localized in AWC cell bodies and axons. **(A)** Images of wild-type L1 animals expressing *unc-116::GFP* and *odr-1p::TagRFP*, which labels both AWCL (left AWC) and AWCR (right AWC). **(B)** Images of a wild-type L1 animal expressing *odr-3p::unc-116::GFP* in AWC neurons. (A, B) Scale bar, 5μm. Anterior to the left and ventral at the bottom.

### *unc-116/kinesin-1* acts non-cell autonomously in the AWC^ON^ neuron to promote the AWC^OFF^ subtype

Previous studies have shown that *unc-104/kinesin-3* acts non-cell autonomously in the pre-AWC^ON^ cell to promote the AWC^OFF^ subtype (Chang *et al*. 2011), which differs from the cell-autonomous role of *tir-1* in promoting the AWC^OFF^ subtype (Chuang and Bargmann 2005). To determine whether *unc-116/kinesin-1* also acts non-cell autonomously to promote the AWC^OFF^ subtype, we analyzed AWC asymmetry phenotypes in mosaic animals overexpressing *unc-116* in one, but not both AWC cells. An extrachromosomal array containing the transgene *odr-3p::unc-116* (expressed in AWC) and *odr-1p::DsRed* (expressed in AWC and AWB) was introduced into animals with the integrated *str-2p::GFP* AWC^ON^ marker. Animals that spontaneously lost the transgene in one AWC cell, but not the other, were identified using the *odr-1p::DsRed* marker. In wild-type animals, the ratio of AWC^ON^, expressing *str-2p::GFP*, between the left and right is approximately equal (Figure 3A, row a) (Troemel *et al*. 1999a), which is consistent with the characteristics of stochastic AWC asymmetry. When the *odr-3p::unc-116* transgene was present in both AWC neurons, the ratio of AWC^ON^ between the left and right cells did not show a statistically significant difference (Figure 3A, row b). In mosaic animals, in which the *odr-3p::unc-116* transgene array was lost in one AWC but retained in the other, 88% of the AWC cells, regardless of left or right, that retained the transgene, acquired the AWC^ON^ subtype (Figure 3A, rows c and d, and 3B). These results suggest that *unc-116/kinesin-1* acts non-cell autonomously in the pre-AWC^ON^ cell to specify the AWC^OFF^ subtype, similar to the non-cell-autonomous role of *unc-104/kinesin-3* and different from the cell-autonomous role *tir-1/SARM1* in promoting AWC^OFF^.

**Figure 3.**
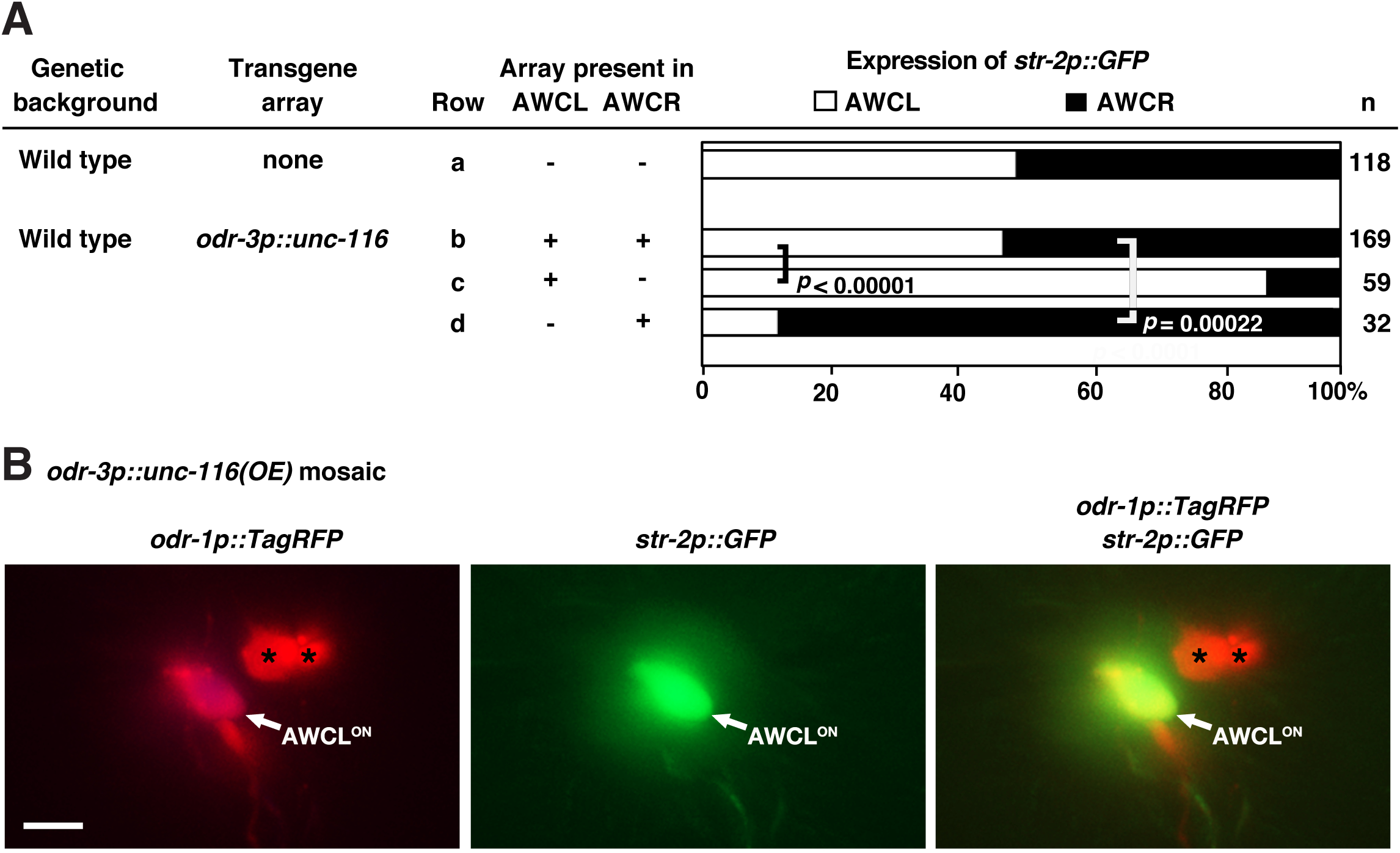
*unc-116* acts non-cell autonomously in the AWC^ON^ neuron to promote the AWC^OFF^ subtype. **(A)** Genetic mosaic analysis of wild-type animals expressing an integrated *str-2p::GFP* transgene and an unstable transgene array, carrying *odr-3p::unc-116* and the mosaic marker *odr-1p::TagRFP*. +, cell retaining the extrachromosomal transgene array; −, cell losing the extrachromosomal transgene array; AWCL, left AWC; AWCR, right AWC. *n*, number of animals scored. Statistical analysis was performed using a *Z*-test. **(B)** Representative image of a mosaic animal expressing the *odr-3p::unc-116* transgene in the AWC^ON^ cell, labeled by the *str-2p::GFP* marker. Asterisks indicate AWB cells. Scale bar, 10μm. Anterior to the left and ventral at the bottom.

### *unc-116/kinesin-1* is required for the localization of TIR-1/SARM1 in the AWC axons

To determine whether *unc-116* regulates the synaptic localization of TIR-1 in AWC cells, the expression pattern of TIR-1::GFP in AWC, expressed from the transgene *odr-1p(-393-170)::tir-1::GFP*, was compared between wild type and *unc-116 (e2310)* mutants. We found that the TIR-1::GFP intensity was significantly reduced in *unc-116(e2310)* mutants compared to the wild type (Figure 4). To rule out the possibility that the *odr-1(-393-170)* promoter was downregulated in *unc-116(e2310)* mutants, GFP expression from the transgene *odr-1p(-393-170)::GFP* was compared between wild type and *unc-116(e2310)* mutants. We found no significant difference in GFP intensity of *odr-1p(-393-170)::GFP* in *unc-116(e2310)* mutants, compared to wild type (Figure S1 A-B). Together, these results suggest that *unc-116/kinesin-1* is required for the synaptic localization of TIR-1/SARM1 in the AWC axons.

**Figure 4.**
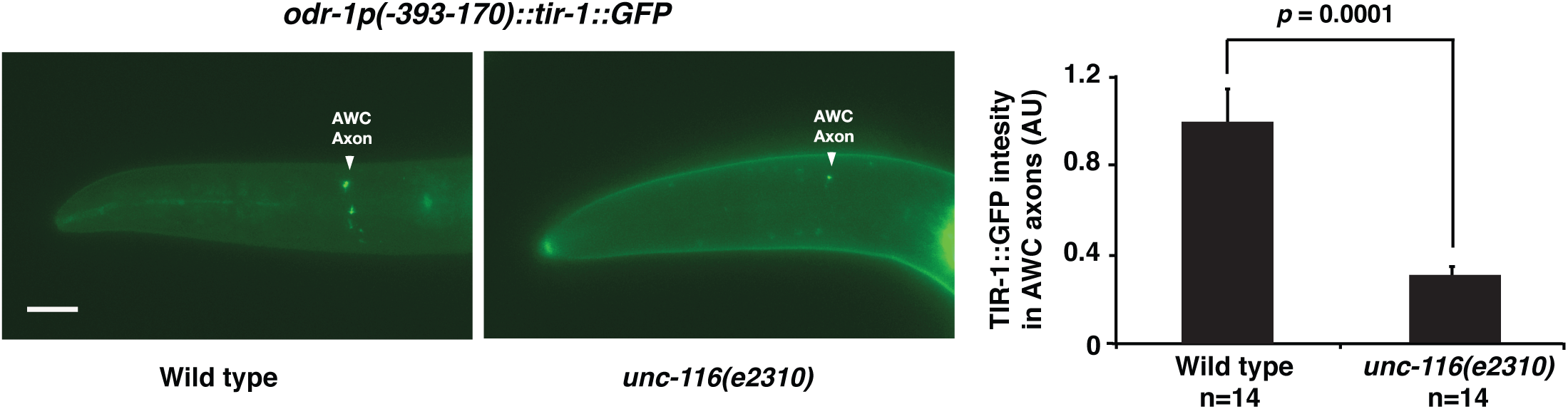
*unc-116/kinesin-1* is required for the localization of TIR-1/SARM1 in the AWC axons. Left panels: Images of wild type and *unc-116(e2310)* mutants expressing *odr-1p(-393-170)::tir-1::GFP* in AWC axons at L1 stage. Scale bar, 10μm. Anterior to the left and ventral at the bottom. Right panel: Quantification of TIR-1::GFP fluorescence intensity in AWC axons. *unc-116(e2310)* mutants displayed a significantly decreased TIR-1::GFP intensity compared to wild type. Statistical analysis was performed using Student’s *t-*test. Error bars, standard error of the mean. AU, arbitrary unit.

### *unc-116/kinesin-1* regulates the dynamic trafficking of TIR-1/SARM1 in the AWC axons

To determine whether the dynamic transport of TIR-1 to the AWC axons is dependent on *unc-116* activity, we analyzed the movement directions and speed of TIR-1::GFP, expressed from the *odr-1p(-393-170)::tir-1::GFP* transgene, along the AWC axons in both wild type and *unc-116(e2310)* mutants. Kymographs were generated from time-lapse images to quantify anterograde and retrograde TIR-1 movement events in AWC axons (Figure 5A).

**Figure 5.**
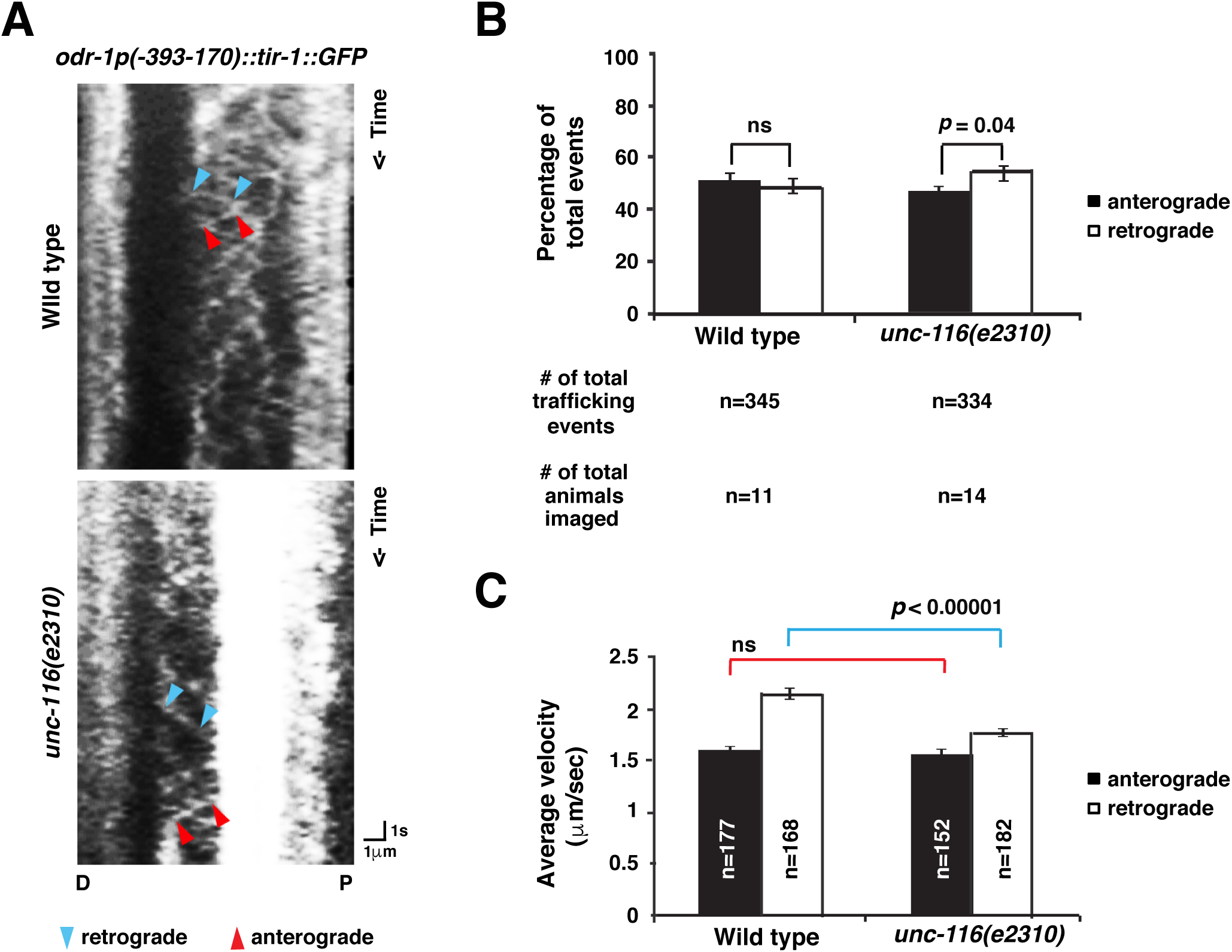
*unc-116/kinesin-1* is required for dynamic trafficking of TIR-1/SARM1 in the AWC axons. **(A)** Representative kymographs of TIR-1::GFP (same transgene as shown in Fig. 4B) movement in AWC axons of wild-type and *unc-116(e2310)* animals at the second and third larval stages. The line within two blue arrowheads represents a retrograde trafficking event from the distal axon toward the proximal (P) axon or the AWC cell body; the line between two red arrowheads represents an anterograde trafficking event away from the cell body down the proximal axon toward the distal (D) axon. **(B)** The percentage of anterograde and retrograde trafficking events. Statistical analysis was performed using a *Z*-test. Error bars, standard error of proportion. ns, not significant. **(C)** The average velocity of anterograde and retrograde trafficking events. Red and blue lines indicate the comparison of anterograde and retrograde velocities, respectively. Statistical analysis was performed using Student’s *t-*test. Error bars, standard error of the mean. ns, not significant.

In wild type, TIR-1::GFP displayed both anterograde (away from the AWC cell body down the proximal axon toward the distal axon) and retrograde (from the distal axon toward the proximal axon or the cell body) movement events with nearly the same ratios (Figure 5B). However, in *unc-116(e2310)* mutants, the ratio of anterograde TIR-1::GFP movement was significantly reduced (Figure 5B). These results are consistent with reduced localization of TIR-1::GFP in the AWC axons in *unc-116(e3210)* mutants. Furthermore, while the average velocity of anterograde movements in the wild type versus *unc-116(e3210)* mutants was not significantly different, the average velocity of retrograde movements was reduced significantly in *unc-116(e2310)* mutants, compared to wild type (Figure 5C). These results suggest that UNC-116 promotes anterograde transport of TIR-1 in the AWC axons by regulating the directionality and relative velocity of movements.

### UNC-116/kinesin-1 is localized adjacent to TIR-1 and colocalized with UNC-104/kinesin-3 in the AWC axons

The punctate pattern of UNC-116/kinesin-1 along AWC axons is similar to that of TIR-1/SARM1 and UNC-104/kinesin-3 (Figure 2B) (Chuang and Bargmann 2005; Chang *et al*. 2011). A previous study showed that UNC-104 and TIR-1 displayed adjacent localization along AWC axons (Chang *et al*. 2011). To determine whether UNC-116 and TIR-1, similar to UNC-104 and TIR-1, also show adjacent localization, UNC-116::GFP and TIR-1::TagRFP were co-expressed in AWC cells from the transgenes *odr-3p::unc-116::GFP* and *odr-3p::tir-1::TagRFP*, respectively. UNC-116::GFP was expressed in a punctate pattern along AWC axons in an adjacent, non-overlapping pattern with TIR-1::TagRFP (Figure 6A), similar to the adjacent localization pattern of UNC-104 and TIR-1 (Chang *et al*. 2011).

**Figure 6.**
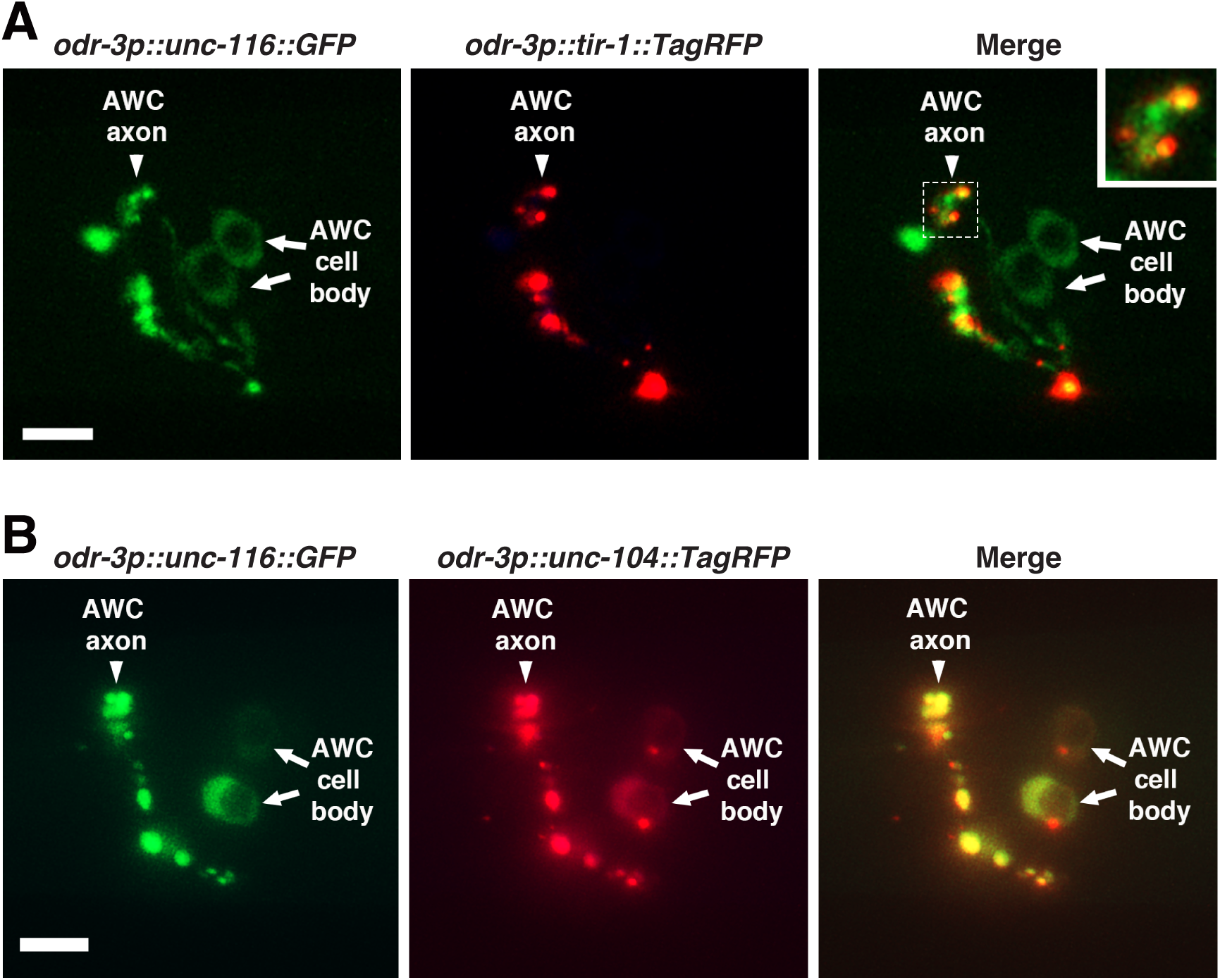
UNC-116/kinesin-1 is localized adjacent to TIR-1 and colocalized with UNC-104/kinesin-3 in the AWC axons. **(A)** Images of a wild-type L1 animal expressing *odr-3p::unc-116::GFP* and *odr-3p::tir-1::TagRFP* in AWC neurons. The inset is magnified by 2-fold. **(B)** Images of a wild-type L1 animal expressing *odr-3p::unc-116::GFP* and *odr-3p::unc-104::TagRFP* in AWC neurons. (A, B) Scale bar, 5μm. Anterior to the left and ventral at the bottom.

Since UNC-116 and UNC-104 have a similar non-cell-autonomous role in promoting AWC^OFF^, we determined whether UNC-116 and UNC-104 colocalize by co-expressing UNC-116::GFP and UNC-104::TagRFP in AWC cells from the transgenes *odr-3p::unc-116::GFP* and *odr-3p::unc-104::TagRFP*, respectively. UNC-116::GFP and UNC-104::TagRFP displayed nearly completely overlapping expression patterns in both AWC cell bodies and axons (Figure 6B).

Together, these results are consistent with the non-cell-autonomous functions of *unc-116/kinesin-1* and *unc-104/kinesin-3* in the pre-AWC^ON^ cell, which trans-synaptically regulate the synaptic localization of TIR-1/SARM1 in the pre-AWC^OFF^ cell to promote the AWC^OFF^ subtype.

## Discussion

Motor proteins in the kinesin/dynein family are key regulators of cellular cargo transport in many cells and have been implicated in neuronal activity (Auer *et al*. 2015; Glomb *et al*. 2023) and cell differentiation (Chang *et al*. 2011; Quesnelle *et al*. 2023). Here, we link the activity of UNC-116, a *C. elegans* ortholog of mouse motor protein KIF-1A of the kinesin-1 family, to the establishment of neuronal asymmetry by the postsynaptic localization of calcium-dependent adaptor protein TIR-1/SARM1. Using a classical genetics approach, we found that the synaptic localization of TIR-1 in AWC neurons is regulated trans-synaptically, not only by UNC-104/kinesin-3, as shown previously (Chang *et al*. 2011), but also by UNC-116/kinesin-1. Our results suggest that both UNC-116/kinesin-1 and UNC-104/kinesin-3 may work together to establish AWC asymmetry in *C. elegans*.

Previous studies have proposed a model in which UNC-104 transports some unknown presynaptic factor(s) in the pre-AWC^ON^ cell, which trans-synaptically regulates the localization of TIR-1 in the pre-AWC^OFF^ cell (Chang *et al*. 2011; HSIEH *et al*. 2014). The non-cell-autonomous requirement of *unc-116* shown in this study suggests that UNC-116/kinesin-1 may also transport the same presynaptic factor(s) as UNC-104/kinesin-3, working together to regulate the synaptic localization of TIR-1/SARM1 in the pre-AWC^OFF^ cell. This model is supported by the nearly overlapping localization of UNC-116 and UNC-104.

Studies have shown that kinesin-1 and kinesin-3 proteins work together, especially in the anterograde transport of the same cellular cargo along microtubules, in which kinesin-1 acts as the dominant motor, though sensitive to roadblocks, while kinesin-3 acts as a co-transporter that is less sensitive to post-translational modifications on microtubules that would otherwise hinder kinesin-1 movement (Lim *et al*. 2017; ArpaĞ *et al*. 2019; Zahavi *et al*. 2021). Furthermore, *in vitro* studies have shown that while kinesin-1 and kinesin-3 can non-redundantly transport cargo in the same direction, they are also able to “take turns,” alternating their activity (Norris *et al*. 2014; Lim *et al*. 2017; ArpaĞ *et al*. 2019). The tradeoff between kinesin-1 and kinesin-3 activity described above may suggest that while UNC-116/kinesin-1 and UNC-104/kinesin-3 have distinct roles in transporting cellular cargo in AWC cells, they may compensate for the loss of one another’s activity. This hypothesis is supported by the wild-type AWC asymmetry displayed by single mutants of both *unc-116* and *unc-104*.

Interestingly, however, we found that *unc-116; unc-104* double mutants did not show any defects in AWC asymmetry, raising the question of whether additional, unidentified motor proteins also work alongside *unc-116* and *unc-104*. Unfortunately, the complete loss of *unc-116/kinesin-1* or *unc-104/kinesin-3* activity is lethal in *C. elegans* (Hall and Hedgecock 1991; Meyerzon *et al*. 2009), limiting our study to the use of partial loss-of-function (or reduction-of-function) alleles, such as *unc-116(e2310)* and *unc-104(e1265),* which may still retain some residual activity of *unc-116* and *unc-104*, respectively. Further studies, using alternative methods, would be beneficial for determining whether UNC-104 (kinesin-1) and UNC-116 (kinesin-3) work together exclusively or alongside other motor proteins to regulate the localization of TIR-1/SARM1 in AWC neurons.

In addition to cell-autonomous functions, kinesins also exhibit non-cell-autonomous functions, where kinesin activity in one cell influences neighboring or related cells through neuronal signaling (Chang *et al*. 2011; Auer *et al*. 2015; Glomb *et al*. 2023; Quesnelle *et al*. 2023). For example, the *C. elegans* kinesin-2 protein Klp-20, which is exclusively expressed in neurons, non-cell autonomously regulates the morphogenesis of epidermal cells (Quesnelle *et al*. 2023). In zebrafish, Kif5aa (kinesin-1) mutants exhibited disrupted presynaptic activity, resulting in non-cell-autonomous upregulation of neutrophin-3 in postsynaptic neurons (Auer *et al*. 2015). Additionally, the *C. elegans* UNC-76 kinesin-1 adaptor protein non-cell autonomously promotes proper axon branching (Glomb *et al*. 2023). Our previous studies have shown that *C. elegans unc-104/kinesin-3* non-cell autonomously regulates the synaptic localization of TIR-1/SARM1 in neighboring neurons for the specification of an olfactory neuron subtype (Chang *et al*. 2011). This study identifies the non-cell-autonomous role of *unc-116/kinesin-1*, alongside *unc-104/kinesin-3*, in regulating the dynamic transport of TIR-1/SARM1 to specify the AWC^OFF^ neuron subtype. The asymmetric differentiation of the AWC neuron pair provides a unique system for shedding light on the mechanisms underlying the non-cell-autonomous functions of kinesin motor proteins in regulating the development of neighboring cells.

## Materials and Methods

### Strains and transgenes

The wild-type *C. elegans* strain is N2, Bristol variety. Strains were cultured by standard methods (Brenner 1974). A list of strains and transgenes is included in Supplemental Materials and Methods.

## Supplemental Materials and Methods

The file includes supplemental strains and transgenes, plasmid construction, supplemental methods, and supplemental references.

## Data availability

All data discussed in the paper will be available to readers.

## Acknowledgments

We thank Paul Mains, Francis McNally, and the *C. elegans* Genetic Center (funded by the NIH Office of Research Infrastructure Programs, P40 OD010440) for strains and reagents. We also thank WormBase. A.K. was supported by a College of Liberal Arts and Sciences Undergraduate Research Initiative (LASURI) Scholarship and a Howard L. Kaufman Scholarship from UIC.

## Author Contributions

Conceptualization, C.-F.C.; Methodology, A.K., J.Y., SY., R.X., and C.-F.C.; Investigation, A.K., J.Y., SY., R.X., and C.-F.C.; Formal Analysis, A.K., J.Y., SY., R.X., and C.-F.C.; Writing, A.K., P.S., and C.-F.C.; Funding Acquisition, A.K. and C.-F.C.; Supervision, C.-F.C.

---

**Figure S1.**
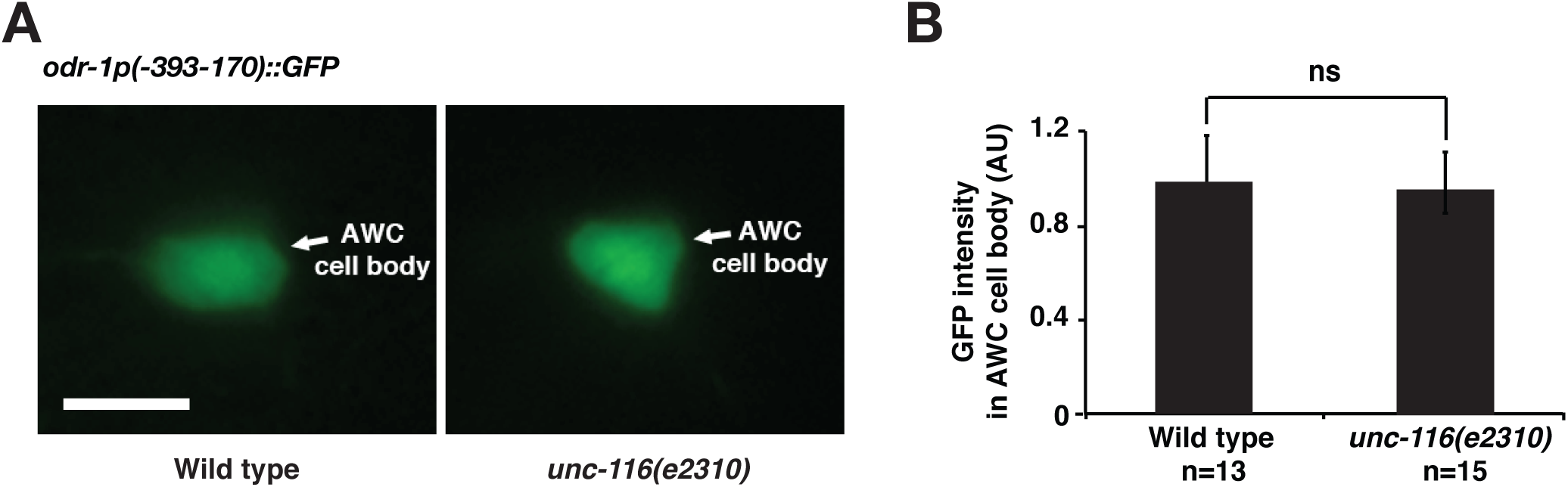
(Related to Figure 4) *unc-116* mutations do not affect the expression levels of *odr-1p (-393-170)* in AWC. **(A)** Representative images of wild type and *unc-116(e2310)* animals expressing *odr-1p(-393-170)::GFP* in the AWC cell body taken at identical exposure times in the first larval stage. Scale bar, 5μm. Anterior to the left and ventral at the bottom. **(B)** Quantification of GFP fluorescence intensity in AWC cell bodies. *unc-116(e2310)* mutants displayed no significant difference in GFP intensity compared to wild type. Statistical analysis was performed using Student’s *t-*test. Error bars, standard error of the mean. AU, arbitrary unit.

## Supplemental Materials and Methods

### Strains and transgenes

Animal protocols approved by the Office of Animal Care and Institutional Biosafety Committees at the University of Illinois Chicago were followed. Hermaphrodites of *C. elegans* were analyzed and imaged.

#### Mutants

*unc-104 (e1265)* II (Hall and Hedgecock 1991)

*tir-1 (ky388)* III (Chuang and Bargmann 2005)

*unc-116 (e2310)* III (Patel *et al*. 1993)

*unc-116 (rh24)* III (Patel *et al*. 1993)

*unc-116 (rh24sb79)* III (Yang *et al*. 2005)

#### Integrated transgenes

| Integrated transgenes | Figures |
| --- | --- |
| *kyIs140* [*str-2p::GFP*; *lin-15(+)*] I (TROEMEL *et al.* 1999) | 1, 3 |
| *vyIs56* [*odr-1p::TagRFP*] III (ALQADAH *et al.* 2016) | 2 |
| *duIs1* [*unc-116::GFP*] (MCNALLY *et al.* 2010) | 2 |

#### Extrachromosomal arrays

| Extrachromosomal arrays | Figures |
| --- | --- |
| *vyEx2496, 2498* [*odr-3p::unc-116 (50 ng/ul)*; *odr-1p::DsRed (15 ng/ul)*; *ofm-1p::DsRed (30 ng/ul)*] | 1, 3 |
| vyEx2666 [*odr-3p::unc-116::GFP* (1 ng/ul); *odr-3p::tir-1::TagRFP* (15ng/ul); *ofm-1p::DsRed* (30 ng/ul)] | 6 |
| vyEx2714 [*odr-3p::unc-116::GFP* (1 ng/ul); *odr-3p::unc-104::TagRFP* (15ng/ul); *ofm-1p::DsRed* (30 ng/ul)] | 2, 6 |
| vyEx2473 [*odr-1p(-393-170)::tir-1::GFP* (7.5 ng/ul); *ofm-1p::DsRed* (30 ng/ul)] | 4, 5 |
| vyEx1609 [*odr-1p(-393-170)::GFP* (50 ng/ul); *ofm-1p::DsRed* (30 ng/ul)] | S1 |

### Plasmid construction

*odr-3p::unc-116* was generated by PCR amplifying a 3290 bp *unc-116* coding region from genomic DNA (Forward primer: GCACTAGCTAGCATGGAGCCGCGGACAG; Reverse primer: CATGCAGGTACCCTTGCTTGAAAAACTTTGATTAGTGAAAAATGGACG), which was then subcloned into a vector containing the *odr-3* promoter and *unc-54* 3’ UTR.

*odr-3p::unc-116::GFP* was made by subcloning the *unc-116 gDNA* from *odr-3p::unc-116 gDNA* into a vector containing the *odr-3* promoter and *GFP* with *unc-54* 3’ UTR.

*odr-3p::tir-1::TagRFP* was constructed by replacing GFP in *odr-3p::tir-1a cDNA::GFP* (Chuang and Bargmann 2005) with TagRFP.

*odr-3p::unc-104::TagRFP* was made by subcloning 4752 bp of *unc-104* cDNA into the pCFJ356 vector (FrØkjÆr-Jensen *et al*. 2012) containing the *odr-3* promoter and *TagRFP* with *unc-54* 3’ UTR.

*odr-1p(-393-170)::tir-1::GFP* was generated by subcloning 2987 bp of *tir-1a* isoform cDNA into a vector containing the *odr-1(-393-170)* promoter and *GFP* with *unc-54* 3’ UTR (Alqadah *et al*. 2015).

### Germline transformation

In brief, a DNA mix was injected into the syncytial gonad of adult hermaphrodites (P_0_) as previously described (Mello and Fire 1995). F_1_ animals expressing the co-injected fluorescent transgenes were identified and cloned (1 animal per plate), and the F_2_ progenies were screened and selected for transgenic lines.

### Genetic mosaic analysis

Animals containing unstable extrachromosomal transgene arrays were passed for at least six generations before performing mosaic analysis, as previously described (Sagasti *et al*. 2001; Vanhoven *et al*. 2006). The co-injection marker *odr-1p::DsRed* (expressed in both AWC neurons) was used to identify the mosaic animals that lose the extrachromosomal transgene in one of the two AWC cells.

### Live imaging of transgenic animals expressing fluorescent proteins

Animals were mounted onto 2% agarose pads and anesthetized with 5 mM sodium azide (Sigma) or 7.5 mM levamisole (Sigma). Z-stack images were obtained using a Zeiss Axio Imager M2 microscope, equipped with a motorized focus drive, a Zeiss objective EC Plan-Neofluar 40x/1.30 Oil DIC M27, a Piston GFP bandpass filter set (41025, Chroma Technology), a TRITC filter set (41002c, Chroma Technology), a Hamamatsu digital camera C11440, and a Zeiss Apotome system. Images were acquired using Zeiss imaging software, ZEN (2012 Blue Edition SP2).

### Quantification of fluorescence intensity

Animals for each set of experiments were imaged using the same exposure time. Fluorescence intensity was measured using the Zeiss imaging software ZEN. In Figure S1, a single focal plane with the brightest GFP fluorescence in the AWC cell body was selected from Z-stack images to compare fluorescence intensity. The quantification analysis was performed by the same individual.

### Time-lapse imaging of protein trafficking

Time-lapse imaging was performed as previously described (Chang *et al*. 2011; Siete *et al*. 2024). Worms in the second larval stage were anesthetized with 7.5 mM tetramisole and mounted onto 2% agarose pads on microscope slides for imaging. We previously showed that 7.5 mM tetramisole, compared to 0.5 mM, 1 mM, and 2 mM tetramisole, did not significantly affect the axonal transport of TIR-1::GFP along the AWC axons (Siete *et al*. 2024). Time-lapse images were acquired for 30 seconds with an exposure time of 300 milliseconds and a speed of 3 frames per second using a Zeiss Axio Imager M2 microscope, equipped with a Zeiss objective EC Plan-Neofluar 63x/1.40 Oil DIC M27, a Piston GFP bandpass filter set (41025, Chroma Technology), a Hamamatsu digital camera C11440, and the Zeiss imaging software ZEN (2012 blue edition SP2). Acquired images were analyzed to generate kymographs using the Fiji software (Schindelin *et al*. 2012). The percentage and velocity of moving events were analyzed using ImageJ software.

